# Heritable intraspecific variation among prey in size and movement interact to shape predation risk and potential natural selection

**DOI:** 10.1101/2024.01.11.575285

**Authors:** Kyle E. Coblentz, Liuqingqing Yang, Arpita Dalal, Miyauna M. N. Incarnato, Dinelka D. Thilakarathne, Cameron Shaw, Ryan Wilson, Francis Biagioli, Kristi L. Montooth, John P. DeLong

## Abstract

1. Predator and prey traits are important determinants of the outcomes of trophic interactions. In turn, the outcomes of trophic interactions shape the evolution of predator and prey traits. How species’ traits are likely to respond to selection from trophic interactions depends crucially on whether and how heritable species’ traits are and their genetic correlations. Of the many traits potentially influencing the outcomes of trophic interactions, body size and movement traits have emerged as universally key traits. Yet, how these traits shape and are shaped by trophic interactions is generally unclear, as few studies have simultaneously measured the impacts of these traits on the outcomes of trophic interactions, their heritability, and their correlations within the same system.
2. We used outcrossed and then clonally propagated lines of the ciliate protist *Paramecium caudatum* from natural populations to examine variation in morphology and movement behavior, the extent to which this variation was heritable, and how intraspecific differences among lines altered *Paramecium* susceptibility to predation by the copepod *Macrocyclops albidus*.
3. We found that the *Paramecium* lines exhibited heritable variation in body size and movement traits. In contrast to expectations from interspecific allometric relationships, body size and movement speed showed little covariance among clonal lines. The proportion of *Paramecium* consumed by copepods was positively associated with *Paramecium* body size and velocity, however, we also found evidence of an interaction such that greater velocities led to greater predation risk for large body-sized paramecia but did not alter predation risk for smaller body-sized paramecia. The proportion of paramecia consumed was not related to copepod body size. These patterns of predation risk and heritable trait variation in paramecia suggest that copepod predation may act as a selective force operating independently on movement and body size and generating the strongest selection against large, high-velocity paramecia.
4. Our results illustrate how the interactions between ecology and genetics can shape the potential natural selection on prey traits through the outcomes of trophic interactions. Further simultaneous measures of predation outcomes, traits, and their quantitative genetic characteristics will provide insights into the evolutionary ecology of species interactions and their eco-evolutionary consequences.

## Introduction

The traits of predators and their prey both shape, and are shaped by, the outcomes of trophic interactions (Abrams, 2000; DeLong, 2021; Pimentel, 1961; Schaffer & Rosenzweig, 1978). Predator and prey traits shape the outcome of trophic interactions because they partially determine the results of each step in the foraging process (DeLong, 2021; Jeschke et al., 2002; Wootton et al., 2023). The outcomes of the foraging process, in turn, shape predator and prey traits because, all else equal, predators with traits that lead to greater predation success will have higher fitness whereas prey with traits that lead to greater avoidance of predation will have higher fitness (Abrams, 2000; DeLong, 2021; Pimentel, 1961; Schaffer & Rosenzweig, 1978). Thus, intraspecific variation in predator and prey traits plays a key ecological and evolutionary role by both determining the outcomes of trophic interactions for individuals while also providing the raw material necessary for predators or prey to evolve in response to selection through trophic interactions.

Many traits are likely to influence the outcomes of trophic interactions such as prey crypsis and defenses and predator foraging behavior (Endler, 1978; Greene, 1986; Tollrian & Harvell, 1999), and intraspecific variation in these traits alters the likelihood of predation between predator and prey individuals. Among the myriad traits that influence the outcome of trophic interactions, predator and prey body sizes and movement behavior have emerged as key traits that universally influence trophic interactions (Aljetlawi et al., 2004; Pawar et al., 2012; Vucic-Pestic et al., 2010; Yodzis & Innes, 1992). Predator and prey body sizes have direct effects on predator-prey interactions by determining, for example, whether a predator can physically consume a given prey item (Paine, 1976), the energetic demand of predators (Kleiber, 1932; Yodzis & Innes, 1992), and the total amount of energy contained in an individual prey (Charnov, 1976; Yodzis & Innes, 1992). The movement behavior of predators and prey also plays a key role in determining the outcomes of trophic interactions because movement behavior is a direct determinant of the encounter rates among predators and prey. In general, greater predator or prey velocities lead to more encounters among predators and prey that, in turn, lead to more opportunities for predation events (Aljetlawi et al., 2004; Pawar et al., 2012).

Furthermore, in many species body size and movement are inextricably linked. Specifically, a common pattern both within and among species is that predator and prey body sizes and movement exhibit a positive allometric relationship in which larger body sizes are associated with greater movement speeds (Cloyed et al., 2021; Hirt et al., 2017). In general, prior studies have suggested that, for a given prey size, increasing predator size should increase feeding rates through greater encounter rates and/or shorter handling times on prey (Coblentz et al., 2023; Pawar et al., 2012; Rall et al., 2012; Uiterwaal & DeLong, 2020). Similarly, for a given predator body size, increasing prey size could lead to higher predator feeding rates by increasing the rates of encounters among predators and prey or decrease feeding rates by lengthening handling times on prey (Coblentz et al., 2023; Pawar et al., 2012; Rall et al., 2012; Uiterwaal & DeLong, 2020). Prior studies also have found unimodal relationships between feeding rates and the ratios of predator to prey sizes (Brose et al., 2008; Kratina et al., 2022; Rall et al., 2012; Vucic-Pestic et al., 2010). However, the studies in which these unimodal relationships occur often consider ranges in predator-prey body size ratios that cover orders of magnitude in variation.

Despite general expectations for how intraspecific variation in body size and movement may influence the outcomes of trophic interactions, whether and how predator and prey body sizes and movement traits might evolve in response to selection due to trophic interactions depends critically on how heritable these traits are. For any quantitative trait to be able to evolve, some proportion of the variation among individuals in that trait must be heritable (Lande & Arnold, 1983; Lush, 1937). Furthermore, how responsive a trait is to selection depends on how heritable that trait is (Lande & Arnold, 1983; Lush, 1937). If a trait is only weakly heritable, its response to selection will be weak for a given amount of phenotypic trait variation within the population, whereas traits with high heritability will respond more strongly to selection. When traits that influence predator-prey interactions are heritable, they can have important ecological consequences. For example, heritability in prey defense traits can generate eco-evolutionary feedbacks generating unique predator-prey cycles (Yoshida et al., 2003) or structure predator communities when predators differ in their susceptibility to prey defenses (Lenhart et al., 2018). Regarding body sizes and movement, previous research has shown that these traits are likely to be heritable to some extent in most organisms with evidence of heritability in both traits being found across a variety of taxa from microbes to vertebrates (Charmantier et al., 2011; Dochtermann et al., 2019; Gervais et al., 2020; Hertel et al., 2020; Mousseau & Roff, 1987; Stirling et al., 2002).

Beyond heritability, potential responses of traits to selection also depend on the genetic correlations among traits (Arnold, 1992; Lande & Arnold, 1983). For example, genetic correlations can constrain evolutionary responses of traits or lead to evolutionary changes in traits that are not under selection if they are correlated with traits that are under selection (Arnold, 1992). One would expect from allometric relationships between body size and movement that these traits should be positively correlated (Cloyed et al., 2021; Hirt et al., 2017). If this is the case, then selection that occurs in the same direction on both traits or directional selection on either trait alone should allow for a directional evolutionary response in both traits (Arnold, 1992; Lande & Arnold, 1983). However, if selection operates in different directions on the two traits, then evolution may be constrained, and the net outcome of selection is likely to be dependent on the quantitative genetic details of the system (Arnold, 1992). If body size and movement traits are not genetically correlated, however, then selection should be able to operate independently on each trait. Thus, different potential genetic correlations and selection pressures due to predation can lead to different expectations for how natural selection is likely to operate on species.

Previous studies on variation across and within species in body sizes and movement and their effects on trophic interactions, their heritability, and potential genetic correlations provide us with a set of potential expectations on the interactions between predator and prey traits, predation, and evolution. Furthermore, these links between genetics, ecology, and evolution are likely to have important consequences for the functioning of predator-prey interactions. However, it is unclear how these processes interact as we are unaware of any studies that simultaneously have examined the heritability and correlations of body size and movement and their impacts on predation. Here we do so by taking advantage of a laboratory predator-prey system consisting of a ciliate protist prey *Paramecium caudatum* and its copepod predator *Macrocyclops albidus*. We examined how genetically diverse, outcrossed and then clonally propagated lines of *P. caudatum* varied in morphological and movement related traits, the extent to which those traits are heritable, and how intraspecific variation in *Paramecium* body size and movement traits and body size in the copepod *M. albidus* are related to *Paramecium* predation risk. From the prior literature outlined above, we first hypothesized that variation in *Paramecium* body size and movement speed would be heritable and positively correlated because these traits have generally been found to be heritable and are commonly positively correlated in inter- and intra-specific comparisons including in other ciliate protists. Second, we hypothesized that *Paramecium* body size and movement speed would be positively correlated with predation rates because we expected greater body sizes and movement rates to lead to greater encounter rates between the paramecia and copepods and we expected that potential decreases in feeding rates due to handling times would be minimal given that copepods are much larger than the paramecia. Third, we hypothesized that copepod body size would be positively correlated with predation rates on the paramecia because the larger copepod size would also increase encounter rates with the paramecia and potentially lower their handling times.

## Materials and Methods

### Study System

We collected our focal species, the ciliate *Paramecium caudatum*, from three sites near Lincoln, Nebraska, USA: Spring Creek Prairie Audubon Center (40°41′24′′N, 96°51′0′′W), Conestoga Lake State Recreation Area (40°45′36′′N, 96°51′0′′W), and Wildwood Lake State Wildlife Management Area (41°2′24′′N, 96°50′24′′W) in June and July of 2023. We focused our collections on shallow, nearshore waters with emergent or floating vegetation. When we found paramecia in the water, we isolated individual cells. We then washed them four times with autoclaved pond water collected from the Spring Creek Prairie site by pipetting the cell in as small of a volume as possible, placing the cell into 1mL of autoclaved pond water, and repeating this process three more times. After washing the cells, we placed the cells alone in separate test tubes. In total, this generated over one hundred isolated lineages. We reared isolated lineages in lettuce media inoculated with bacteria collected from the Spring Creek Prairie site. We made lettuce media using 15g of broken up organic romaine lettuce autoclaved in 1L of filtered pond water with 0.7g of ground dried autoclaved pond mud to supply rare elements. We maintained the bacterial flora in this media by transferring inoculated media into new jars with uninoculated media every other day or so.

Conjugation is the sexual stage of paramecia and involves meiosis followed by genetic exchange between individuals of different mating types. To generate outcrossed lines from the isolated lineages of *Paramecium*, in August 2023, we combined cells from all isolates into 100mm Petri dishes to promote conjugation. Cells began conjugating within a day, and we collected adjoined conjugates and isolated them into new tubes. As *Paramecium* exconjugates (individual cells post conjugation) are genetically identical (Ahsan et al., 2022; Bell, 1989; Hiwatashi, 2001), individuals descended from the conjugating pair are clones with the potential to be genetically different from clonal lines that descend from other exconjugant pairs through both recombination and genetic differences among conjugating individuals. We established 132 of these outcrossed and then clonally propagated lines and maintained them on lettuce media.

The focal predator in our foraging experiment, the copepod *Macrocyclops albidus*, also originated from the Spring Creek Prairie Audubon Center in June through August, 2023. We used a combination of wild-collected adult and lab-reared individuals in foraging trials. For the lab-reared individuals, we isolated gravid *M. albidus* in a single Petri dish with *P. caudatum* provided *ad libitium* as food. Eggs hatched and grew through stages, and we collected new adults from these stocks for the trials.

We reared all paramecia and copepod stocks at room temperature (23°C).

### Video phenotyping

To examine whether and how the outcrossed *Paramecium* lines differed in morphological and movement traits, we phenotyped cells from videos. Twenty-four hours prior to taking videos of the *Paramecium*, we placed cells from each of the outcrossed lines into fresh bacterized media in new test tubes at room temperature to create common-garden conditions. For each of the outcrossed lines, we washed approximately 20 *Paramecium* cells three times in 1mL of 0.2µm filtered autoclaved pond water. We then placed the *Paramecium* onto a Petri dish in 0.1mL of filtered autoclaved pond water and covered the drop with a deep-well projection slide cover (Carolina Deep-Well Slides, Model: 60-3730 60-3730E). Immediately after placing the slide cover over the *Paramecium*, we took a 25s video of the *Paramecium* using a stereo microscope (Leica M165C) outfitted with a camera (Leica DMC 4500). As some of the outcrossed lines did not have enough cells available on the day of video phenotyping, we ended up with videos of 126 of the 133 outcrossed lines.

To extract morphological and movement data from the videos, we used the R package Bemovi (BEhavior and MOrphology from VIdeo; (Pennekamp et al., 2015)). Bemovi uses particle tracking software to identify and track individual cells in videos and then extracts information on the morphology and movement of cells. For each video, we ran the Bemovi analysis and then averaged the extracted data across all the cells within an outcrossed line to get a single set of average morphological and movement measurements. The number of identified particle tracks used to obtain the averages ranged from N=9-36 identified particles per outcrossed line after data processing and filtering.

### Foraging Experiment

We housed 58 copepods alone within deep-well projection slides (Carolina Deep-Well Slides, Model: 60-3730 60-3730E) of 2.4cm diameter and 1.4mm depth with approximately 0.8mL of filtered autoclaved pond water. Prior to each foraging trial, we starved the copepods for 24 hours to standardize hunger levels. We used only non-gravid copepods. For copepods that became gravid during the experiment, we fed them paramecia daily until they dropped their eggs and could be used again after a 24-hour starvation period. Before each trial, we washed each copepod twice by removing 0.6mL of water from around the copepod and replacing it with filtered pond water. For each trial, we also washed 40 paramecia three times using 1mL filtered pond water. After adding the 40 washed paramecia to the deep-well projection slide arenas, we placed the slide arenas in an incubator (Percival E30BC8) at 25°C. After 30 minutes of foraging, we removed the slide arenas and counted the number of remaining *Paramecium* cells under a stereo microscope. Given the short period of the foraging trials, we did not perform control experiments to account for mortality or cell divisions during the trial. The amount of growth should be minimal since the trial duration is approximately the time it takes for a *P. caudatum* cell to divide once it shows signs of binary fission and we did not use cells that showed signs of division (Hinrichs, 1928). Furthermore, mortality from sources other than copepods is unlikely as we have never observed mortality in healthy paramecium cells in such a short period in autoclaved pond water (Authors, *Personal Observation*).

In total, we conducted 4-7 replicate foraging trials with different *M. albidus* individuals for each of 38 randomly chosen clonal lines of *Paramecium*. Across these trials, the same copepod was never used with the same outcrossed line more than once and was never used more than once in a day. Overall, individual copepods were used in 2-8 total foraging trials. This led to a total of 230 foraging trials.

### Copepod morphology measurements

We photographed 57 of the 58 *M. albidus* used in the foraging trials using a stereo microscope (Leica M165C) outfitted with a camera (Leica DMC 4500) and measured their lengths and widths in millimeters. The one copepod that was not photographed died during the experiment before photographs were taken, and its foraging trials were removed from the data prior to analysis.

### Replication statement for foraging trials

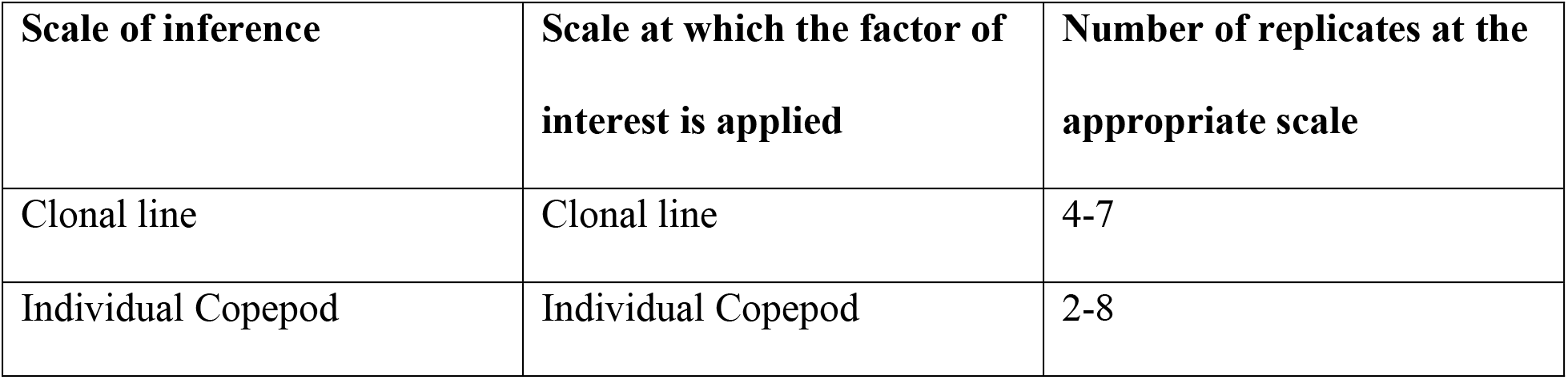

### Statistical Analyses

#### Analysis of Phenotype Data

The Bemovi analysis of the videos provided a suite of potential variables to describe the morphological and movement phenotypes of the *Paramecium* outcrossed lines. To reduce the dimensionality of the phenotypes, we first excluded variables that are given by Bemovi but have an unclear meaning in terms of the *Paramecium* phenotypes such as the ‘grey-ness’ of the paramecia in the videos. Next, we examined a correlation matrix of the remaining variables using the medians across individuals within outcrossed lines to determine which variables were highly correlated with one another and thus may be providing the same or very similar information (see Supplementary Online Material (SOM) 1 for the correlation matrix). For the variables that were highly correlated, we chose the most easily interpretable variable to include as a measure of the phenotype. For example, net speed and net displacement were highly correlated and so we chose to only include net displacement as a descriptor of the *Paramecium* phenotype. After this process we were left with the following variables describing the *Paramecium* phenotypes: length, width, aspect ratio (the ratio of length to width), mean turning angle, standard deviation of the turning angle, gross speed, net displacement, and standard deviation of the gross speed (for definitions of each of these variables see SOM 2). After selecting these variables to describe the *Paramecium* phenotypes, we then further reduced the dimensionality by performing a Principal Components Analysis across the selected variables after centering and standardizing each variable.

We also estimated trait heritability as a measure of the genetic contribution to the variation of each of the selected variables we used to describe the *Paramecium* phenotypes. The amount of measured trait variation among clonal lines can be used to estimate broad-sense trait heritability *H^2^* (Lynch & Walsh, 1998). To estimate the heritability of traits using clonal lines, one can use an Analysis of Variance (ANOVA) with the trait of interest measured for each cell in the videos as the response and the genotype (or clonal line, in this case) as a fixed effect (Lynch & Walsh, 1998). The broad-sense heritability is then estimated as the amount of variation explained by genotype divided by the total variation (Lynch & Walsh, 1998), with the caveat that maternal effects are not factored out of this estimate of *H^2^*.

#### Analysis of Copepod Foraging Data

To analyze the copepod foraging data, we used a generalized linear mixed effects model. To allow for over/under-dispersion in the data, we modeled the response (the proportion of paramecia consumed in the foraging trial) as beta-binomially distributed using a logit link function. To account for the lack of independence due to the repeated use of outcrossed lines and individual copepods, we included random intercepts for *Paramecium* outcrossed line and individual copepod. As our questions were about how *Paramecium* phenotypes and copepod size influenced the proportion of paramecia consumed, we included the first and second principal component analysis axes from the analysis of the *Paramecium* phenotypes, their interaction, and copepod length as fixed effects. Copepod length and width were correlated and using width rather than length had no qualitative effect on our results (SOM 3). We performed the regression in a Bayesian framework using the R package ‘brms’ (Bürkner, 2017). For model details including priors and an assessment of model fit, see Supplemental Material SOM 4. Last, we used predictions from the generalized linear mixed effects model to visualize a fitness surface in which we defined fitness as the predicted proportion of paramecia surviving after the foraging trial dependent on the first and second principal component analysis axes.

All analyses were performed using R v. 4.3.1 (R Core Team, 2023).

## Results

### Paramecium Phenotypes

The first two axes of the Principal Components Analysis explained 60.2% of the total variation in the mean phenotypes across all the lines (Figure 1; see Supplemental Material SOM 5 for the full PCA results). The first axis was positively associated with measures of speed and displacement (i.e. gross speed, its standard deviation, and net displacement) and the aspect ratio of the paramecia and negatively associated with the mean turning angle of the paramecia. The second axis was positively associated with *Paramecium* size (length, width) and had a slight negative association with the standard deviation in turning angle. Thus, we interpret the first axis as representing movement speed and lack of turning in the paramecia and the second axis as a measure of size. We also note that the fact that size and speed loaded on separate axes reflects that size and speed were not strongly correlated among the outcrossed lines (the absolute values of the correlations between the size and movement traits considered ranged from 0.02-0.36 with a mean of 0.14; SOM 1).

**Figure 1.**
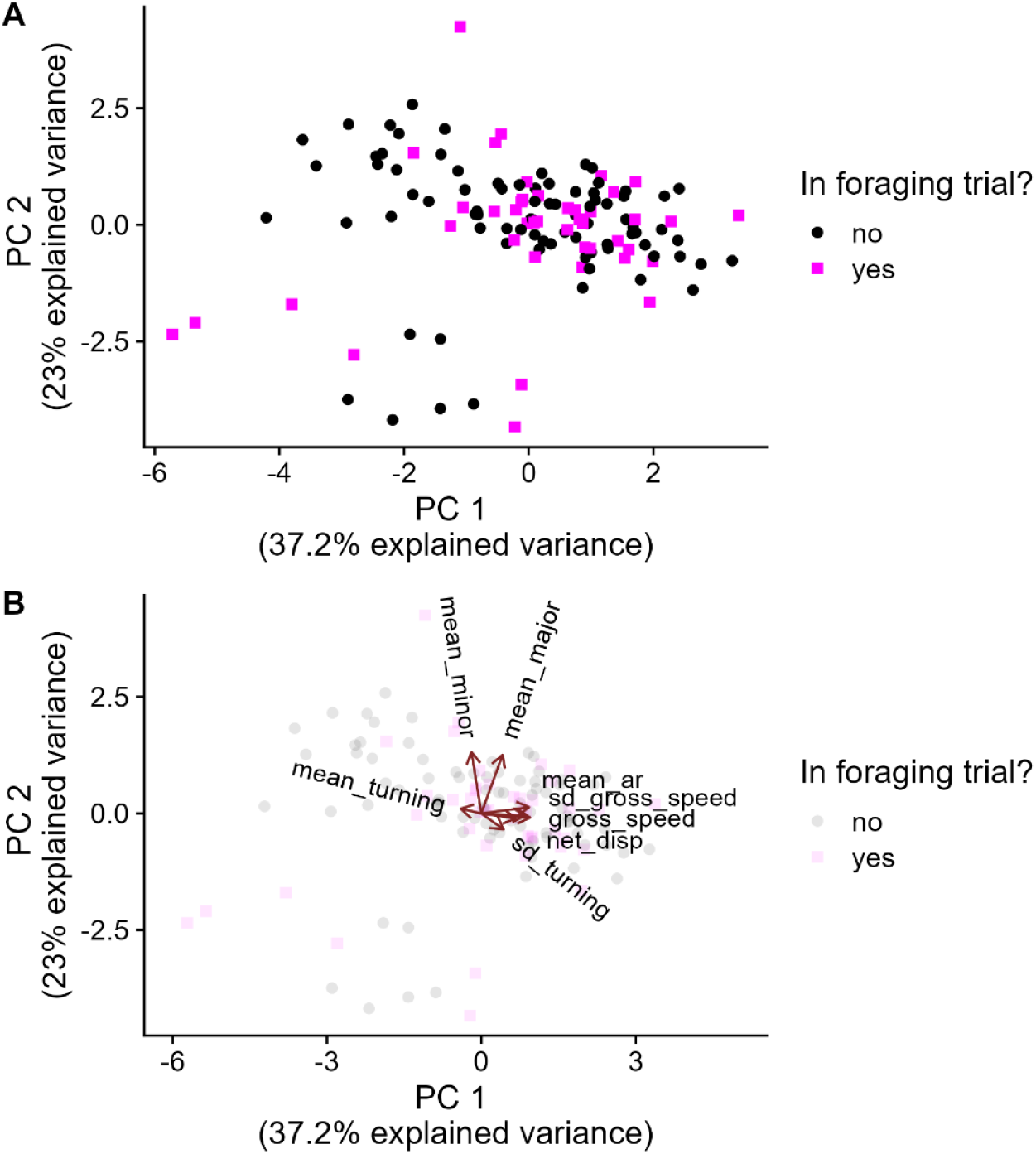
The first two components of a principal components analysis of the mean phenotypes of *Paramecium caudatum* outcrossed lines derived from the analysis of videos explained 60.2% of the total variation (A). The first principal component was positively associated with the mean aspect ratio of the paramecia (mean_ar) and several speed related phenotypes (e.g. gross speed (gross_speed) and net displacement (net_disp)) and negatively associated with the mean turning angle (mean_turning; B). The second principal component was positively associated with the mean length and width of the paramecia (mean_major, mean_minor; B). For definitions of each of the phenotypic traits included in the principal components analysis see Supplementary Online Material SOM2 and SOM5. The black circles in A and B represent the 88 outcrossed lines that were phenotyped but not included in the foraging trials whereas the magenta squares represent the 38 outcrossed lines that were phenotyped and included in the foraging trials.

Estimates of the broad-sense heritability of the morphological and movement traits of the paramecia ranged from 0.14 to 0.73 (Table 1). The estimates of heritability of the size traits were 0.64 for length, 0.59 for width, and 0.44 for the aspect ratio. The heritability for movement related traits of the paramecia were lower on average than the morphological traits but had greater variation ranging from 0.14 for net displacement to 0.73 for gross speed (Table 1).

**Table 1.** Summary of quantitative genetics results examining the heritability of *Paramecium caudatum* traits.

| **Trait** | **Trait Sum of Squares** | **Total Variance** | **Broad-sense Heritability** |
| --- | --- | --- | --- |
| Major Axis (Length) | $1.4 \times 10^6$ | $2.2 \times 10^6$ | 0.64 |
| Minor Axis (Width) | $2.0 \times 10^5$ | $3.4 \times 10^5$ | 0.59 |
| Aspect Ratio<br>(Length/Width) | 133.3 | 239.3 | 0.44 |
| Mean Turning | 2.1 | 13.9 | 0.15 |
| sd Turning | 42.9 | 111.9 | 0.38 |
| Gross Speed | $3.7 \times 10^8$ | $5.1 \times 10^8$ | 0.73 |
| Net Displacement | $2.2 \times 10^9$ | $15.3 \times 10^9$ | 0.14 |
| sd Gross Speed | $4 \times 10^7$ | $6.5 \times 10^7$ | 0.62 |

### Prey and predator traits and foraging rates

We estimated a positive effect of size, speed, and their interaction on the proportion of *Paramecium* cells offered that were eaten by copepods (Figure 2; Table 2). Together these interactive effects were such that small paramecium had relatively constant risk of predation by copepods whereas large paramecium had greater risk of predation when they were fast (Figure 2, Figure 3). We found no statistically clear relationship between copepod size and the proportion of *Paramecium* offered that were eaten by copepods (Table 2, SOM 6).

**Figure 2.**
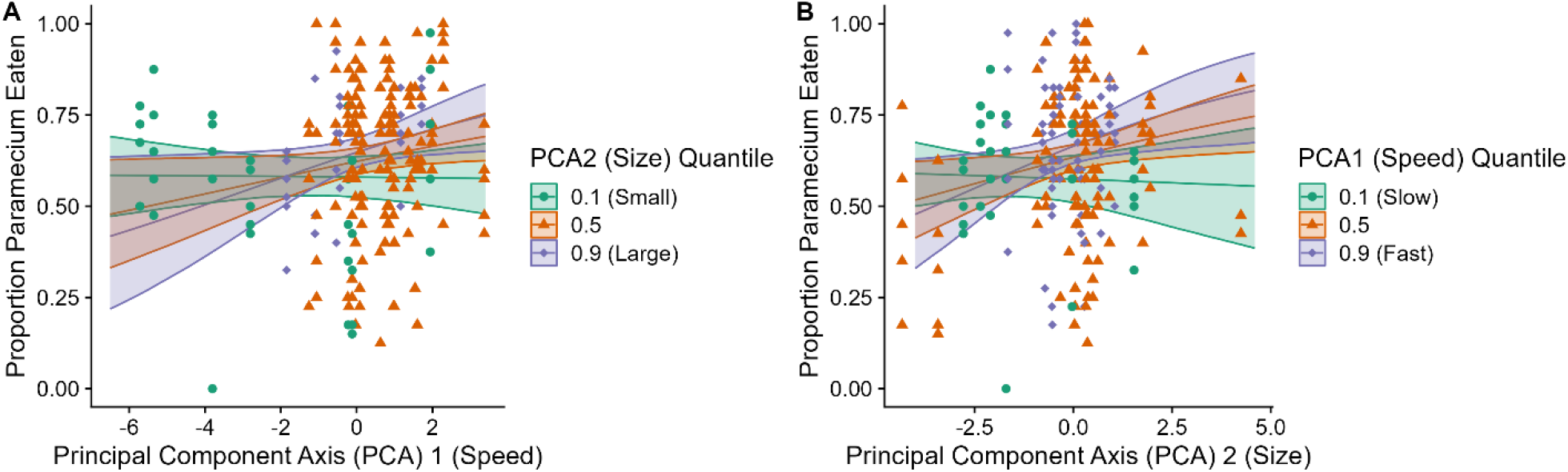
The proportion of paramecium eaten per trial increased with paramecium speed (PCA1) and paramecium size (PCA2). However, there was evidence of an interaction in which the effect of speed was dependent on size such that larger, faster individuals were at greater predation risk whereas small individuals had similar risk regardless of speed. The differently colored lines and shaded areas represent the means and 90% Credible intervals for the relationship between the principal component values and the mean proportion of *Paramecium* consumed by copepods for different quantiles of the principal component not on the x-axis (PCA2 in panel A and PCA1 in panel B). Points are colored such that the green dots are for outcrossed lines from the 0.15 quantile or less of the principal component not on the x-axis, purple dots are for outcrossed lines from the 0.85 quantile or greater of the principal component not on the x-axis, and the rest of the points are orange.

**Figure 3.**
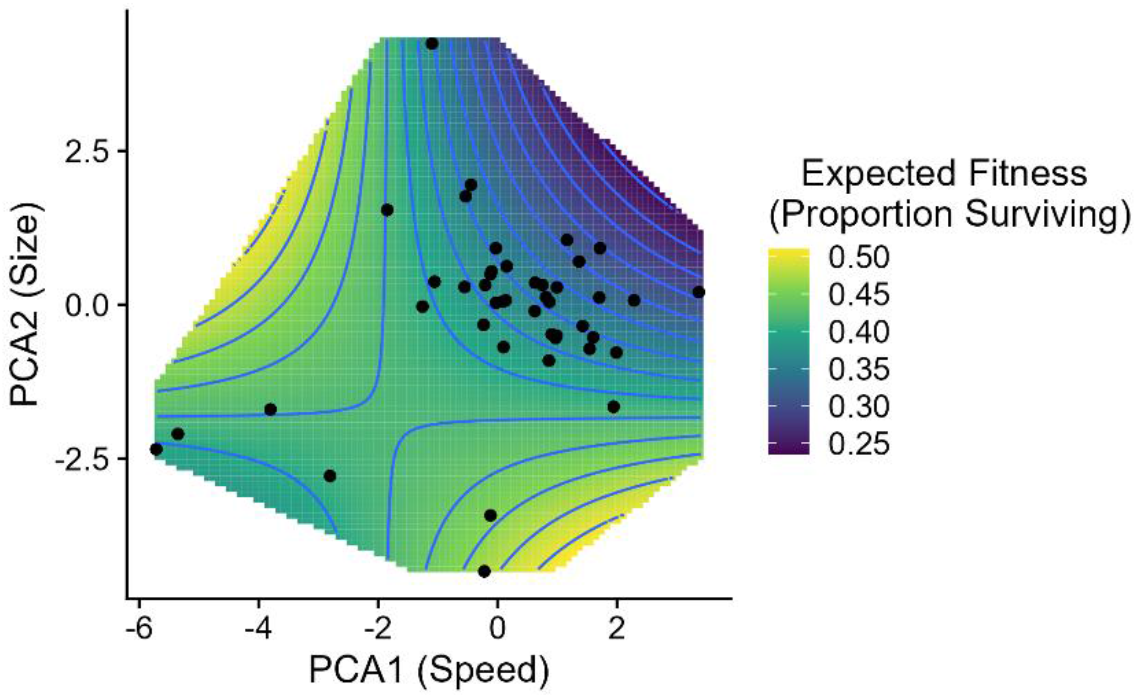
A predicted fitness surface using the expected proportion of paramecia surviving the copepod foraging trials shows that the expected fitness is lowest for paramecia that are both fast (high PCA1 value) and large (a high PCA2 value). Points denote the principal component values for the outcrossed lines included in the foraging experiments, the cut out areas of the fitness surface are those in which there were no outcrossed lines with those combinations of PCA1 and PCA2 values, and the blue lines are contours.

**Table 2.**
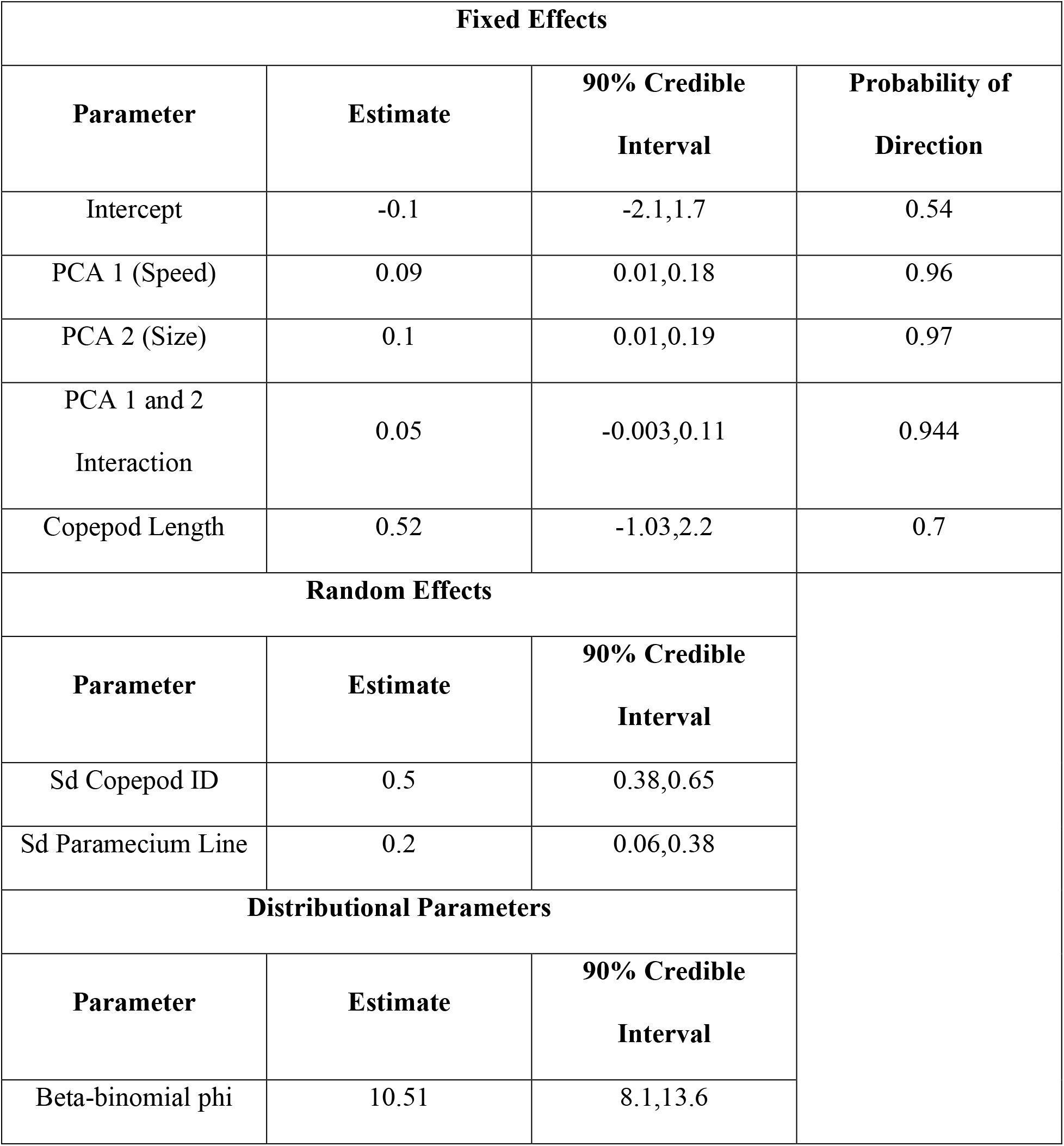
Generalized Linear Mixed Effects Model (GLMM) results examining the relationships between *Paramecium caudatum* traits (PCA 1 and 2) and their interaction and copepod size with the proportion of *P. caudatum* consumed by copepods in feeding trials. Sd in the table stands for ‘standard deviation’.

## Discussion

Intraspecific trait variation plays a critical role in the ecology and evolution of predator-prey interactions. First, the traits of predators and their prey determine the outcomes of trophic interactions and, thus, determine the strengths of predator-prey interactions and their ecological consequences (DeLong, 2021; Wootton et al., 2023). Second, if the trait variation determining the outcomes of trophic interactions is heritable, this variation provides the raw material for selection and the evolution and co-evolution of predator-prey interactions (Abrams, 2000; DeLong, 2021; Pimentel, 1961). Using outcrossed and then clonally propagated lines of *Paramecium caudatum*, we measured the structure and heritability of intraspecific phenotypic variation in two key sets of traits for determining the outcomes of predator-prey interactions – body size and movement -- and determined how these traits were related to the predation risk of paramecia by copepods. Overall, our study revealed some clear differences between hypotheses generated from previous studies of interspecific patterns in movement and morphological traits and their relationships with predation and our results. Furthermore, by simultaneously examining the structure and heritability of variation in *Paramecium* movement and body size traits along with their relationships to predator risk, our analyses also revealed how selection through copepod foraging might operate on *Paramecium* phenotypes.

Studies of allometric scaling have generally shown that species’ movement speeds increase with body size across a wide variety of organisms (e.g. Cloyed et al., 2021; Hirt et al., 2017). Although these studies have largely focused on multicellular organisms, positive relationships between body size and speed have also been found within species in another ciliate protist (Pennekamp et al., 2019). Thus, we hypothesized that movement speed and body size would be correlated with one another across our outcrossed lines of *Paramecium*. Our analysis of the *Paramecium* phenotypes, however, showed that variation in speed and body size had largely low correlations. Instead, the morphological trait that was most positively correlated with speed was the aspect ratio of the paramecia. A previous study on flagellate protists that also showed no relationships with body size and speed, however, may provide a potential explanation (Nielsen & Kiørboe, 2021). In their study, Nielsen and Kiørboe (2021) found that flagellates with smaller widths and flagellar characteristics that generated greater force had increased speeds. These results suggest that the relationship between the aspect ratio of the paramecia and swimming speed may be due to a combination of a decreased radius of the cell coupled with greater ability to generate force for a given width by having a greater number of cilia along the length of the cell. Regardless of the cause of the general independence between size and speed, its existence plays a potentially important role in how paramecia might respond to selection from copepod foraging. Specifically, the independence of size and speed suggests a general lack of genetic correlation between the two sets of traits that can potentially allow the paramecia to separately respond to selection on each set of traits (Lande & Arnold, 1983). Coupled with the observed relationship that predation risk from copepods was highest on fast and large paramecia, the independence of speed and size suggests that selection due to copepod predation could operate to reduce size, speed, or both with similar fitness results.

Many studies have shown how differences among individuals can lead to differences in interactions with other species including predation risk (e.g. Cuthbert et al., 2020; Morgan et al., 2016; Pretorius et al., 2019). Although many of the quantitative traits considered in these studies are likely to be heritable, this assumption is rarely tested because studies often use ontogenetic differences between individuals, for example, in body size, to examine effects of intraspecific variation or do not take the extra step to determine whether differences have a genetic basis. In our study, there is both ontogenetic variation within outcrossed lines due to differences among individuals in time since cell division and variation due to genetic differences among outcrossed lines. In general, our analyses show that the *Paramecium* traits we considered show significant heritability despite ontogenetic variation. We also find that some traits are more heritable than others. For example, all of the size related traits we examined showed a moderate amount of heritability, whereas the movement-related traits showed greater variation in heritability. Previous studies have found that morphological traits, in general, have greater heritabilities than behavioral or physiological traits (Dochtermann et al., 2019; Mousseau & Roff, 1987; Stirling et al., 2002). Our results are consistent with these meta-analyses despite the fact that they were largely based on multicellular organisms and biased towards vertebrates. Given the high heritability estimates for gross speed and its standard deviation that exceed those of the morphological traits, we also hypothesize that there may be a tight link between these movement traits and ciliary morphology or density (Funfak et al., 2014; Osterman & Vilfan, 2011). Overall, the general heritability of morphological and movement traits suggests that although certain traits may be able to respond more readily to selection, the paramecia should be capable of evolving in both size and movement related traits.

Predator and prey body sizes and velocities in cross-species comparisons show clear relationships with predator feeding rates (Coblentz et al., 2023; Pawar et al., 2012; Uiterwaal & DeLong, 2020; Vucic-Pestic et al., 2010). In general, these studies show that predator feeding rates increase with increasing predator size and with higher movement velocities in either species. Further, cross-species allometries between size and velocities suggest higher feeding rates of predators on larger prey due to greater encounter rates (although this could be counteracted by longer handling times in some systems). These patterns led us to hypothesize that: 1) predation risk for the paramecia would increase with *Paramecium* body size and velocity, and 2) predation risk for the paramecia would increase with copepod body size. We found partial support for the hypothesis that *Paramecium* predation risk would increase with body size and velocity. However, we also found evidence of an interaction through which predation risk was highest for large, fast paramecia and was nearly constant for small paramecia regardless of speed. We hypothesize that the reason for this is that smaller paramecia regardless of their encounter rates with copepods may be harder to detect and capture than larger paramecia, leading to similar predation rates on smaller paramecia regardless of their velocity. On the other hand, larger paramecia may be easier to detect and capture, leading to a dependence of predation risk on velocities and encounter rates. These hypotheses are supported by studies of the mechanosensory mechanisms of prey detection in copepods that show larger prey have a greater detection distance than smaller prey (Almeda et al., 2018; Jonsson & Tiselius, 1990; Kiørboe & Visser, 1999). An alternative hypothesis from optimal foraging theory is that the copepods fed more so on larger paramecia due to their higher energy content and ate more of the faster, large paramecia due to higher encounter rates (Charnov, 1976). In general, a reduction in protist size in response to predation has been noted in several predation experiments with protists as prey (Fyda et al., 2005; Griffiths et al., 2018; Kratina et al., 2010; terHorst et al., 2010). Furthermore, although some previous studies have also found unimodal relationships between predation rates and prey size, an analysis of our model residuals showed no evidence of a unimodal relationship (SOM 7). We suspect that there is no unimodal relationship in our study because studies that do show unimodal relationships between predation rates and prey sizes typically span orders of magnitude in predator-prey body mass ratios whereas predator-prey length ratios in our system only ranged from 5 to 13 (Kratina et al., 2022; Rall et al., 2012; Vucic-Pestic et al., 2010). Additional experiments may be able to tease apart the causes of the interaction between paramecia size and speed in determining *Paramecium* predation risk, but our main takeaway is that copepod foraging generates correlational selection – selection on a trait that depends of the value of another trait -- on the paramecia in size and movement (Brodie III, 1992; Lande & Arnold, 1983).

In contrast to *Paramecium* size, we found no evidence of an effect of copepod size on *Paramecium* predation risk. We hypothesize that this may be due to the large body size difference between paramecia and copepods and the relative range of size differences observed in the paramecia versus in the copepods. Again, studies examining the effects of predator and prey body sizes on predator feeding rates, both inter- and intra-specifically, generally measure these effects across orders of magnitude in variation of sizes or predator-prey body size ratios (Coblentz et al., 2023; Englund et al., 2011; Pawar et al., 2012; Rall et al., 2012; Uiterwaal & DeLong, 2020; Vucic-Pestic et al., 2010). In our study, mean paramecium lengths among the outcrossed lines used in the experiment ranged from 95-233µm whereas copepod size ranged from 0.9-1.4mm. It is possible that over a larger range of copepod sizes we would have found an effect of copepod size, but that feeding rates on paramecia are generally similar in the size range that occurs among adult copepods of this species. Despite the lack of an effect of copepod body size on *Paramecium* predation risk, our statistical model suggested that there was substantial variation among copepods through the random effect of individual copepod. As the model showed no statistically clear effect of copepod size and copepod hunger was standardized, variation among copepods may have been due to some uncontrolled factor such as age or behavioral differences among individual copepods (Toscano et al., 2016; Toscano & Griffen, 2014). Although our inclusion of copepod identity as a random effect in our statistical model may have masked the effect of copepod size on the proportion of paramecia consumed, we believe this is unlikely as there was no evident relationship between proportion of paramecium consumed and copepod size (SOM 6). Thus, our results on copepod body size provide an example of a case in which intraspecific variation in predator and prey traits need not match those predicted from cross-species comparisons as has been found in some other predation studies (e.g. DiFiore & Stier, 2023; Gallagher et al., 2016; Gibert et al., 2017).

Altogether, the patterns of heritability and correlations of the paramecia traits coupled with the potential correlational selection imposed by copepod foraging suggest that: 1) there are a variety of morphological and movement trait combinations that lead to similar fitness values, and 2) these areas of similar fitness should be readily accessible by paramecium populations. This suggests that there is no single adaptive peak in terms of paramecium morphology and movement in regard to copepod predation. Rather, there is a broad fitness plateau with similarly high fitness values for small paramecium regardless of their speed and for large, slow paramecium (Figure 3). Although this fitness plateau may exist in the presence of copepod predation, *Paramecium* morphology and movement are likely to have important effects on many other *Paramecium* functions such as competitive ability, dispersal, and their own feeding rates (Gibert et al., 2017; Pennekamp et al., 2019; Tan et al., 2021). In turn, these additional ecological effects of morphology and movement may lead to different fitness landscapes and patterns of selection on the paramecia. Nevertheless, strong copepod predation may act as a filter on *Paramecium* morphology and movement narrowing which combinations of size and speed can lead to high fitness. This reflects the overall grand challenge of understanding how the genetic structure of predator and prey traits interact with multiple sources of selection to shape predator and prey traits, the resultant strengths of those predator-prey interactions given predator and prey traits, and their consequences for populations and communities.

## Conclusions

Understanding the reciprocal relationships between the traits of predators and prey and the outcomes of their interactions is key to understanding the functional ecology and evolution of predator-prey interactions (Abrams, 2000; DeLong, 2021; Pimentel, 1961; Schaffer & Rosenzweig, 1978). To do so requires simultaneous knowledge of how traits influence the outcomes of trophic interactions and the underlying genetics of the traits involved. Using a laboratory system, we were able to determine the heritabilities and correlations of key morphological and movement traits of a prey species and their relationships to predation risk. Our results revealed several mismatches between expectations from cross-species allometric relationships and intraspecific patterns in our prey while also revealing the potential for predators to impose correlational selection on uncorrelated traits. These patterns suggest an ability of the prey to adapt to predation in a diversity of ways that lead to similar fitness outcomes and call for a better reconciliation between patterns of inter- and intraspecific variation. We hope that our study inspires future work integrating quantitative genetics and functional predator ecology to provide a more synthetic understanding of the eco-evolutionary processes determining the outcomes of predator-prey interactions and their ecological and evolutionary consequences.

## Author Contributions

KEC, KLM, and JPD designed the study with input of the other authors, all authors performed the study, KEC performed the statistical analyses and led the writing of the manuscript, all authors helped to write the manuscript or contributed to revisions.

## Statement on inclusion

All authors are currently based in the region where the study was conducted.

## Acknowledgements

This work was funded by an NSF Organismal Responses to Climate Change Grant (ORCC- 2307464) to KLM and JPD. AD was supported by Fulbright-Kalam Climate Fellowship for Postdoctoral Research (Award No. 2867 FNPDR /2022). LY was supported by an NSF Graduate Research Fellowship.

## Data Availability Statement

On acceptance, all data and code for the analysis of the data will be permanently archived as a Github repository on Zenodo.

## References

1. Abrams, P. A. (2000). The Evolution of Predator-Prey Interactions: Theory and Evidence. Annual Review of Ecology, Evolution, and Systematics, 31(Volume 31, 2000), 79–105. 10.1146/annurev.ecolsys.31.1.79

2. Ahsan, R., Blanche, W., & Katz, L. A. (2022). Macronuclear development in ciliates, with a focus on nuclear architecture. Journal of Eukaryotic Microbiology, 69(5), e12898. 10.1111/jeu.12898

3. Aljetlawi, A. A., Sparrevik, E., & Leonardsson, K. (2004). Prey–predator size-dependent functional response: Derivation and rescaling to the real world. Journal of Animal Ecology, 73(2), 239–252. 10.1111/j.0021-8790.2004.00800.x

4. Almeda, R., van Someren Gréve, H., & Kiørboe, T. (2018). Prey perception mechanism determines maximum clearance rates of planktonic copepods. Limnology and Oceanography, 63(6), 2695– 2707. 10.1002/lno.10969

5. Arnold, S. J. (1992). Constraints on Phenotypic Evolution. The American Naturalist, 140, S85–S107. 10.1086/285398

6. Bell, G. (1989). *Sex and Death in Protozoa: The History of Obsession*. Cambridge University Press. 10.1017/CBO9780511525704

7. Brodie III, E. D. (1992). Correlational Selection for Color Pattern and Antipredator Behavior in the Garter Snake Thamnophis Ordinoides. Evolution, 46(5), 1284–1298. 10.1111/j.1558-5646.1992.tb01124.x

8. Brose, U., Ehnes, R. B., Rall, B. C., Vucic-Pestic, O., Berlow, E. L., & Scheu, S. (2008). Foraging theory predicts predator–prey energy fluxes. Journal of Animal Ecology, 77(5), 1072–1078. 10.1111/j.1365-2656.2008.01408.x

9. Bürkner, P.-C. (2017). brms: An R Package for Bayesian Multilevel Models Using Stan. Journal of Statistical Software, 80(1), Article 1. 10.18637/jss.v080.i01

10. Charmantier, A., Buoro, M., Gimenez, O., & Weimerskirch, H. (2011). Heritability of short-scale natal dispersal in a large-scale foraging bird, the wandering albatross. Journal of Evolutionary Biology, 24(7), 1487–1496. 10.1111/j.1420-9101.2011.02281.x

11. Charnov, E. L. (1976). Optimal Foraging: Attack Strategy of a Mantid. The American Naturalist, 110(971), 141–151. 10.1086/283054

12. Cloyed, C. S., Grady, J. M., Savage, V. M., Uyeda, J. C., & Dell, A. I. (2021). The allometry of locomotion. Ecology, 102(7), e03369. 10.1002/ecy.3369

13. Coblentz, K. E., Novak, M., & DeLong, J. P. (2023). Predator feeding rates may often be unsaturated under typical prey densities. Ecology Letters, 26(2), 302–312. 10.1111/ele.14151

14. Cuthbert, R. N., Wasserman, R. J., Dalu, T., Kaiser, H., Weyl, O. L. F., Dick, J. T. A., Sentis, A., McCoy, M. W., & Alexander, M. E. (2020). Influence of intra- and interspecific variation in predator–prey body size ratios on trophic interaction strengths. Ecology and Evolution, 10(12), 5946–5962. 10.1002/ece3.6332

15. DeLong, J. P. (2021). Predator Ecology: Evolutionary Ecology of the Functional Response. Oxford University Press.

16. DiFiore, B. P., & Stier, A. C. (2023). Variation in body size drives spatial and temporal variation in lobster– urchin interaction strength. Journal of Animal Ecology, 92(5), 1075–1088. 10.1111/1365-2656.13918

17. Dochtermann, N. A., Schwab, T., Anderson Berdal, M., Dalos, J., & Royauté, R. (2019). The Heritability of Behavior: A Meta-analysis. Journal of Heredity, 110(4), 403–410. 10.1093/jhered/esz023

18. Endler, J. A. (1978). A Predator’s View of Animal Color Patterns. In M. K. Hecht, W. C. Steere, & B. Wallace (Eds.), Evolutionary Biology (pp. 319–364). Springer US. 10.1007/978-1-4615-6956-5_5

19. Englund, G., Öhlund, G., Hein, C. L., & Diehl, S. (2011). Temperature dependence of the functional response. Ecology Letters, 14(9), 914–921. 10.1111/j.1461-0248.2011.01661.x

20. Funfak, A., Fisch, C., Motaal, H. T. A., Diener, J., Combettes, L., Baroud, C. N., & Dupuis-Williams, P. (2014). Paramecium swimming and ciliary beating patterns: A study on four RNA interference mutations. Integrative Biology, 7(1), 90–100. 10.1039/C4IB00181H

21. Fyda, J., Warren, A., & Wolinńska, J. (2005). An investigation of predator-induced defence responses in ciliated protozoa. Journal of Natural History, 39(18), 1431–1442. 10.1080/00222930400004396

22. Gallagher, A. J., Brandl, S. J., & Stier, A. C. (2016). Intraspecific variation in body size does not alter the effects of mesopredators on prey. Royal Society Open Science, 3(12), 160414. 10.1098/rsos.160414

23. Gervais, L., Hewison, A. J. M., Morellet, N., Bernard, M., Merlet, J., Cargnelutti, B., Chaval, Y., Pujol, B., & Quéméré, E. (2020). Pedigree-free quantitative genetic approach provides evidence for heritability of movement tactics in wild roe deer. Journal of Evolutionary Biology, 33(5), 595– 607. 10.1111/jeb.13594

24. Gibert, J. P., Allen, R. L., Hruska III, R. J., & DeLong, J. P. (2017). The ecological consequences of environmentally induced phenotypic changes. Ecology Letters, 20(8), 997–1003. 10.1111/ele.12797

25. Greene, C. H. (1986). Patterns of Prey Selection: Implications of Predator Foraging Tactics. The American Naturalist, 128(6), 824–839. 10.1086/284608

26. Griffiths, J. I., Petchey, O. L., Pennekamp, F., & Childs, D. Z. (2018). Linking intraspecific trait variation to community abundance dynamics improves ecological predictability by revealing a growth– defence trade-off. Functional Ecology, 32(2), 496–508. 10.1111/1365-2435.12997

27. Hertel, A. G., Niemelä, P. T., Dingemanse, N. J., & Mueller, T. (2020). A guide for studying among-individual behavioral variation from movement data in the wild. Movement Ecology, 8(1), 30. 10.1186/s40462-020-00216-8

28. Hinrichs, M. A. (1928). Ultra-Violet Radiation and Division in Paramecium caudatum. Physiological Zoology, 1(3), 394–415. 10.1086/physzool.1.3.30151054

29. Hirt, M. R., Lauermann, T., Brose, U., Noldus, L. P. J. J., & Dell, A. I. (2017). The little things that run: A general scaling of invertebrate exploratory speed with body mass. Ecology, 98(11), 2751–2757. 10.1002/ecy.2006

30. Hiwatashi, K. (2001). My Paramecium Study, Retrospect and Prospect. Zoological Science, 18(1), 1–4. 10.2108/zsj.18.1

31. Jeschke, J. M., Kopp, M., & Tollrian, R. (2002). Predator Functional Responses: Discriminating Between Handling and Digesting Prey. Ecological Monographs, 72(1), 95–112. 10.1890/0012-9615(2002)072[0095:PFRDBH]2.0.CO;2

32. Jonsson, P., & Tiselius, P. (1990). Feeding behaviour, prey detection and capture efficiency of the copepod Acartia tonsa feeding on planktonic ciliates. Marine Ecology Progress Series, 60, 35–44. 10.3354/meps060035

33. Kiørboe, T., & Visser, A. W. (1999). Predator and prey perception in copepods due to hydromechanical signals. Marine Ecology Progress Series, 179, 81–95. 10.3354/meps179081

34. Kleiber, M. (1932). Body size and metabolism. Hilgardia, 6(11), 315–353.

35. Kratina, P., Hammill, E., & Anholt, B. R. (2010). Stronger inducible defences enhance persistence of intraguild prey. Journal of Animal Ecology, 79(5), 993–999.

36. Kratina, P., Rosenbaum, B., Gallo, B., Horas, E. L., & O’Gorman, E. J. (2022). The Combined Effects of Warming and Body Size on the Stability of Predator-Prey Interactions. Frontiers in Ecology and Evolution, 9. 10.3389/fevo.2021.772078

37. Lande, R., & Arnold, S. J. (1983). The Measurement of Selection on Correlated Characters. Evolution, 37(6), 1210–1226. 10.2307/2408842

38. Lenhart, P. A., Jackson, K. A., & White, J. A. (2018). Heritable variation in prey defence provides refuge for subdominant predators. Proceedings of the Royal Society B: Biological Sciences, 285(1879), 20180523. 10.1098/rspb.2018.0523

39. Lush, J. L. (1937). Animal Breeding Plans. Collegiate Press, Incorporated.

40. Lynch, M., & Walsh, B. (1998). Genetics and Analysis of Quantitative Traits. Oxford University Press.

41. Morgan, S. G., Gravem, S. A., Lipus, A. C., Grabiel, M., & Miner, B. G. (2016). Trait-mediated indirect interactions among residents of rocky shore tidepools. Marine Ecology Progress Series, 552, 31–46. 10.3354/meps11766

42. Mousseau, T. A., & Roff, D. A. (1987). Natural selection and the heritability of fitness components. Heredity, 59(2), 181–197. 10.1038/hdy.1987.113

43. Nielsen, L. T., & Kiørboe, T. (2021). Foraging trade-offs, flagellar arrangements, and flow architecture of planktonic protists. Proceedings of the National Academy of Sciences, 118(3), e2009930118. 10.1073/pnas.2009930118

44. Osterman, N., & Vilfan, A. (2011). Finding the ciliary beating pattern with optimal efficiency. Proceedings of the National Academy of Sciences, 108(38), 15727–15732. 10.1073/pnas.1107889108

45. Paine, R. T. (1976). Size-Limited Predation: An Observational and Experimental Approach with the Mytilus-Pisaster Interaction. Ecology, 57(5), 858–873. 10.2307/1941053

46. Pawar, S., Dell, A. I., & Van M. Savage. (2012). Dimensionality of consumer search space drives trophic interaction strengths. Nature, 486(7404), Article 7404. 10.1038/nature11131

47. Pennekamp, F., Clobert, J., & Schtickzelle, N. (2019). The interplay between movement, morphology and dispersal in Tetrahymena ciliates. PeerJ, 7, e8197. 10.7717/peerj.8197

48. Pennekamp, F., Schtickzelle, N., & Petchey, O. L. (2015). BEMOVI, software for extracting behavior and morphology from videos, illustrated with analyses of microbes. Ecology and Evolution, 5(13), 2584–2595. 10.1002/ece3.1529

49. Pimentel, D. (1961). Animal Population Regulation by the Genetic Feed-Back Mechanism. The American Naturalist, 95(881), 65–79. 10.1086/282160

50. Pretorius, J. D., Lichtenstein, J. L. L., Eliason, E. J., Stier, A. C., & Pruitt, J. N. (2019). Predator-induced selection on urchin activity level depends on urchin body size. Ethology, 125(10), 716–723. 10.1111/eth.12924

51. R Core Team. (2023). R: A Language and Environment for Statistical Computing. R Foundation for Statistical Computing. https://www.R-project.org/

52. Rall, B. C., Brose, U., Hartvig, M., Kalinkat, G., Schwarzmüller, F., Vucic-Pestic, O., & Petchey, O. L. (2012). Universal temperature and body-mass scaling of feeding rates. Philosophical Transactions of the Royal Society B: Biological Sciences, 367(1605), 2923–2934. 10.1098/rstb.2012.0242

53. Schaffer, W. M., & Rosenzweig, M. L. (1978). Homage to the red queen. I. Coevolution of predators and their victims. Theoretical Population Biology, 14(1), 135–157. 10.1016/0040-5809(78)90008-4

54. Stirling, D. G., Réale, D., & Roff, D. A. (2002). Selection, structure and the heritability of behaviour. Journal of Evolutionary Biology, 15(2), 277–289. 10.1046/j.1420-9101.2002.00389.x

55. Tan, H., Hirst, A. G., Atkinson, D., & Kratina, P. (2021). Body size and shape responses to warming and resource competition. Functional Ecology, 35(7), 1460–1469. 10.1111/1365-2435.13789

56. terHorst, C. P., Miller, T. E., & Levitan, D. R. (2010). Evolution of prey in ecological time reduces the effect size of predators in experimental microcosms. Ecology, 91(3), 629–636. 10.1890/09-1481.1

57. Tollrian, R., & Harvell, C. D. (1999). *The Ecology and Evolution of Inducible Defenses*. Princeton University Press. 10.2307/j.ctv1ddd1cn

58. Toscano, B. J., Gownaris, N. J., Heerhartz, S. M., & Monaco, C. J. (2016). Personality, foraging behavior and specialization: Integrating behavioral and food web ecology at the individual level. Oecologia, 182(1), 55–69. 10.1007/s00442-016-3648-8

59. Toscano, B. J., & Griffen, B. D. (2014). Trait-mediated functional responses: Predator behavioural type mediates prey consumption. Journal of Animal Ecology, 83(6), 1469–1477. 10.1111/1365-2656.12236

60. Uiterwaal, S. F., & DeLong, J. P. (2020). Functional responses are maximized at intermediate temperatures. Ecology, 101(4), e02975. 10.1002/ecy.2975

61. Vucic-Pestic, O., Rall, B. C., Kalinkat, G., & Brose, U. (2010). Allometric functional response model: Body masses constrain interaction strengths. Journal of Animal Ecology, 79(1), 249–256. 10.1111/j.1365-2656.2009.01622.x

62. Wootton, K. L., Curtsdotter, A., Roslin, T., Bommarco, R., & Jonsson, T. (2023). Towards a modular theory of trophic interactions. Functional Ecology, 37(1), 26–43. 10.1111/1365-2435.13954

63. Yodzis, P., & Innes, S. (1992). Body Size and Consumer-Resource Dynamics. The American Naturalist, 139(6), 1151–1175. 10.1086/285380

64. Yoshida, T., Jones, L. E., Ellner, S. P., Fussmann, G. F., & Hairston, N. G. (2003). Rapid evolution drives ecological dynamics in a predator–prey system. Nature, 424(6946), Article 6946. 10.1038/nature01767

