## Supplemental Information for "Heritable intraspecific variation among prey in size and movement interact to shape predation risk and potential natural selection"

**Contents** **Page**


| | |
|---|---|
| **SOM 1: *Parmecium* trait correlation matrix** | **2** |
| **SOM 2: Table of *Paramecium* trait definitions** | **3** |
| **SOM 3: Copepod length and width correlation** | **4** |
| **SOM 4: Details of Generalized Linear Mixed Effects Model** | **5** |
| **SOM 5: Principal Component Analysis Results of *Paramecium* phenotypic traits** | **8** |
| **SOM 6: Copepod Size and *Paramecium* Predation Risk** | **9** |
| **SOM 7: Model Residuals and Nonlinearity in Size-Predation Risk Relationships** | **10** |
| **Literature Cited** | **11** |

**SOM 1: *Parmecium* trait correlation matrix**

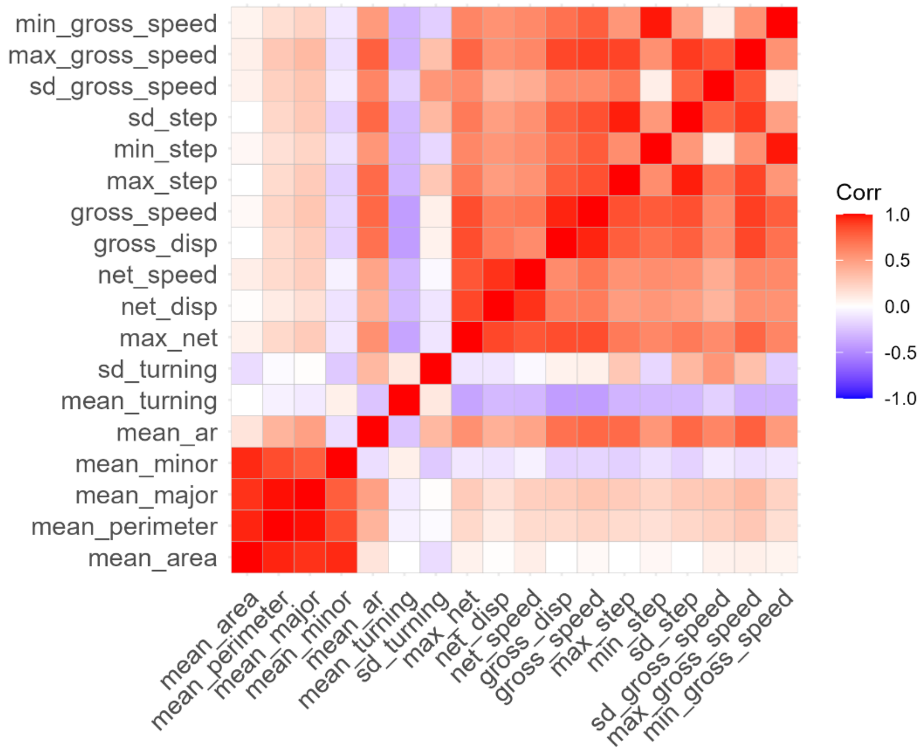


**Figure SOM 1.1.** A correlation matrix of the mean phenotypic traits of outcrossed lines of *Paramecium caudatum*.

### SOM 2: Table of *Paramecium* trait definitions

**Table SOM 2.1.** Traits from the analysis of videos of *Paramecium caudatum* outcrossed line phenotypes and their definitions following Pennekamp et al. (2015).

| Trait | Definition | Abbreviation |
|---|---|---|
| Mean Major Axis | The mean length of a cell across the entire video from fitted ellipses in each frame | mean_major |
| Mean Minor Axis | The mean width of a cell across the entire video from fitted ellipses in each frame | mean_minor |
| Mean Area | The mean 2D area of the cell across the entire video calculated using the Mean Major and Minor Axes | mean_area |
| Mean Perimeter | The mean perimeter of the cell across the entire video | mean_perimeter |
| Mean Aspect Ratio | The mean length/width of a cell across the entire video from fitted ellipses in each frame | mean_ar |
| Mean Turning Angle | The mean of relative changes in cell direction compared across adjacent frames of the video | mean_turning |
| Standard Deviation Turning Angle | The standard deviation of relative changes in cell direction compared across adjacent frames of the video | sd_turning |
| Gross Speed | The mean step lengths taken by the cell across frames divided by the total time between the frames | gross_speed |
| Gross Displacement | The total sum of step lengths taken by the cell across all frames | gross_disp |
| Maximum Net Displacement | The maximum straight line distance a cell appeared in the video from its starting point | max_net |
| Net Displacement | Straight-line distance between the starting point of the cell and the finishing point of the cell | net_disp |
| Net Speed | The net displacement of the cell divided by the time length of time the cell was tracked | net_speed |
| Maximum Step Size | The maximum distance moved by a cell between two frames | max_step |
| Minimum Step Size | The minimum distance moved by a cell between two frames | min_step |
| Standard Deviation of Step Size | The standard deviation of the distance moved by cells between adjacent frames (i.e. step size) | sd_step |
| Maximum Gross Speed | The maximum gross speed achieved by the cell | max_gross_speed |
| Minimum Gross Speed | The minimum gross speed achieved by the cell | min_gross_speed |
| Standard Deviation of Gross Speed | The standard deviation of the gross speed | sd_gross_speed |

**SOM 3: Copepod length and width correlation**

The lengths and widths of the individual copepods *Macrocyclops albidus* used in our foraging experiments were positively correlated (Pearson Correlation Coefficient = 0.67; 95% Confidence Interval = (0.5, 0.79), $p < 2.2 \times 10^{-16}$) and using copepod width rather than length in our regression model had no qualitative effects on our results.

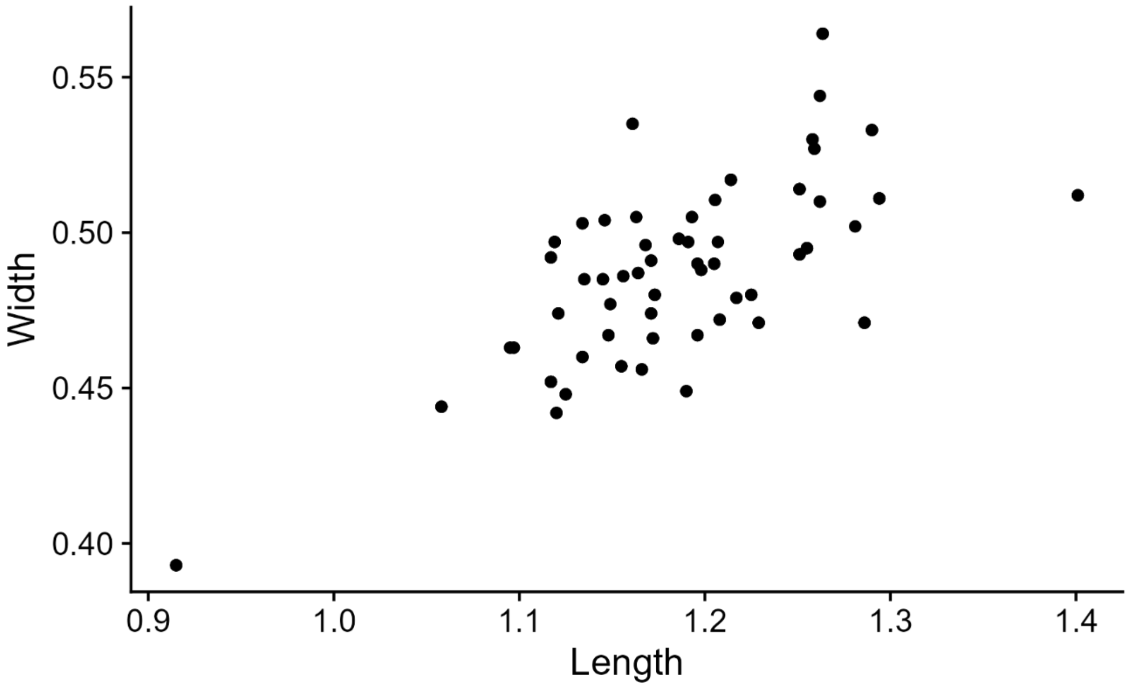


**Figure SOM 3.1.** The length and width of the copepods (in mm) used in our foraging experiments were positively correlated.

### SOM 4: Details of Generalized Linear Mixed Effects Model

To examine the relationships between *Paramecium* morphological and movement traits and copepod size on the risk of predation of *Paramecium* by copepods, we used a Generalized Linear Mixed Effects Model (GLMM). Our response variable was the proportion of the 40 *Paramecium* offered to the copepods that were eaten over the course of 30 minutes which we modeled as beta-binomially distributed to allow for potential over/under-dispersion using a logit link function. As explanatory variables, we included the first two principal components of the Principal Component Analysis on the *Paramecium* morphological and movement data, their interaction, and the length of the copepods. To account for non-independence due to using the same copepods and *Paramecium* outcrossed lines multiple times during the experiment, we included random intercepts for individual copepod and *Paramecium* outcrossed line. We fit the GLMM in a Bayesian framework with Cauchy(location = 0, scale = 2) priors on the intercept, the regression coefficients, and the random intercepts. We also placed a half-Cauchy prior with scale equal to 2 on the standard deviation of the random intercepts. Last, we used the default Gamma(shape = 0.01, scale = 0.01) prior on the 'phi' parameter of the beta-binomial distribution. To approximate the posterior distribution of the parameters, we used 2,000 samples each from four Hamiltonian Monte Carlo chains after 2,000 warmup iterations. Below, we include figures showing that the Hamiltonian Monte Carlo chains for each parameter converged and were well mixed (Figure S4.1; also evidenced by $\hat{R}$ values close to one for all parameters). We also include a posterior predictive check showing that density of predicted data from our model matches the observed distribution of data well (Figure S4.2). The model was fit using the R package 'brms' (Bürkner, 2017; R Core Team, 2023)

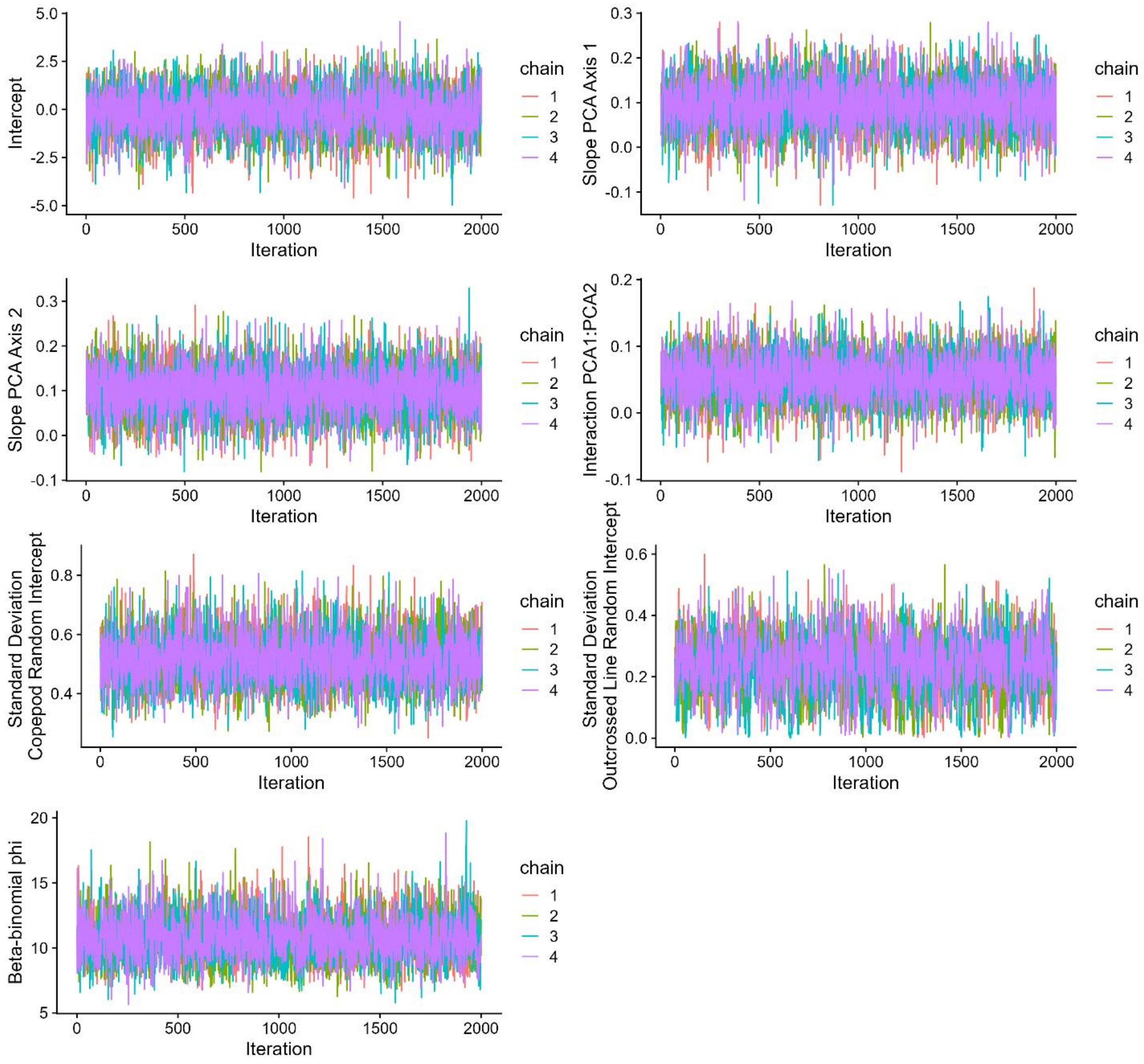


**Figure S4.1.** Trace plots for each parameter of the Bayesian GLMM showing that the Hamiltonian Monte Carlo chains are well-mixed and have converged.

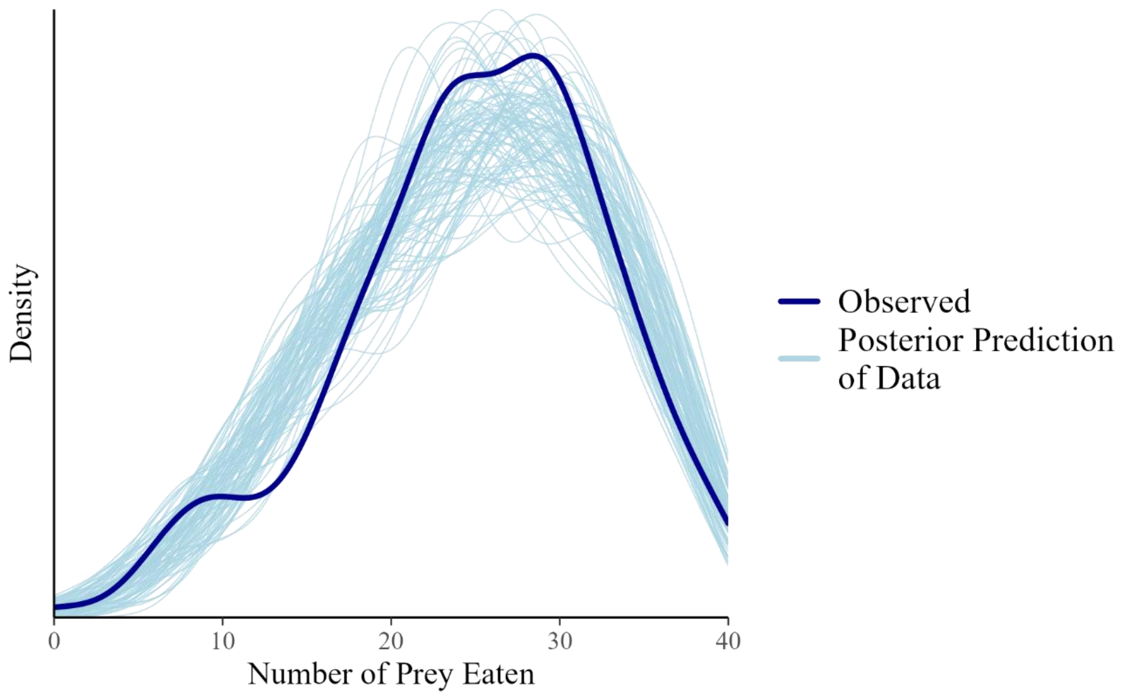


**Figure S4.2.** Density plots of observed data (number of *Paramecium* eaten) as the bold line and predicted data from the approximated posterior distribution showing that the model predicts data that looks similar to the observed data suggesting an overall good fit of the model to the data.

### SOM 5: Principal Component Analysis Results of *Paramecium* phenotypic traits

**Table S5.1.** Results from a Principal Component Analysis of the mean morphological and movement data across 126 outcrossed lines of *Paramecium caudatum* measured from video analysis on 20 cells per outcrossed line.

| Principal Component | PC1 | PC2 | PC3 | PC4 | PC5 | PC6 | PC7 | PC8 |
|---|---|---|---|---|---|---|---|---|
| Mean Major Axis | 0.22 | 0.67 | -0.08 | 0.003 | 0.12 | 0.12 | 0.06 | 0.68 |
| Mean Minor Axis | -0.1 | 0.71 | 0.003 | -0.08 | -0.29 | 0.01 | -0.2 | -0.6 |
| Aspect Ratio | 0.49 | 0.07 | -0.14 | 0.1 | 0.59 | 0.18 | 0.4 | -0.42 |
| Mean Turning | -0.21 | 0.05 | -0.5 | 0.80 | -0.02 | -0.22 | -0.007 | -0.006 |
| Standard Deviation Turning | 0.23 | -0.18 | -0.64 | -0.19 | -0.25 | 0.54 | -0.32 | -0.002 |
| Gross Speed | 0.51 | -0.05 | 0.21 | 0.20 | 0.18 | -0.27 | -0.74 | -0.01 |
| Net Displacement | 0.36 | -0.06 | 0.42 | 0.46 | -0.54 | 0.38 | 0.23 | -0.003 |
| Standard Deviation Gross Speed | 0.46 | -0.03 | -0.28 | -0.23 | -0.42 | -0.62 | 0.3 | 0.007 |
| Proportion of Variance Explained | 0.37 | 0.23 | 0.18 | 0.09 | 0.06 | 0.04 | 0.02 | 0.002 |

**SOM 6: Copepod Size and *Paramecium* Predation Risk**

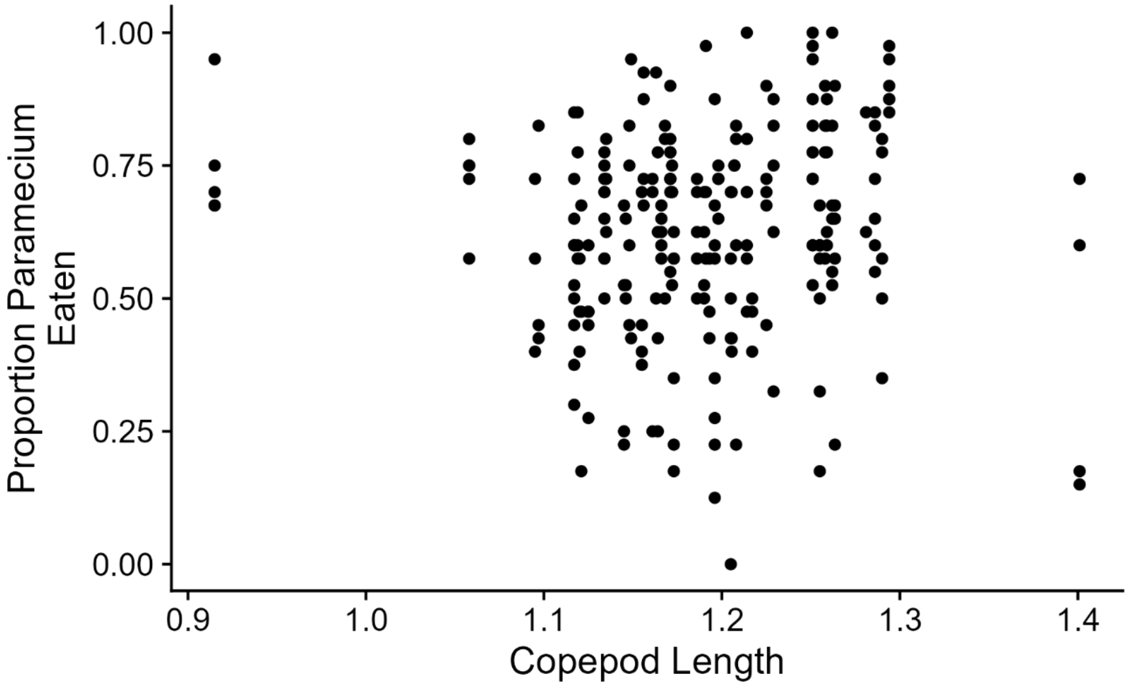


**Figure S6.1.** Copepod size (length in mm) showed no statistically clear relationship with the number of *Paramecium* consumed in foraging trials.

**SOM 7: Model Residuals and Nonlinearity in Size-Predation Risk Relationships**

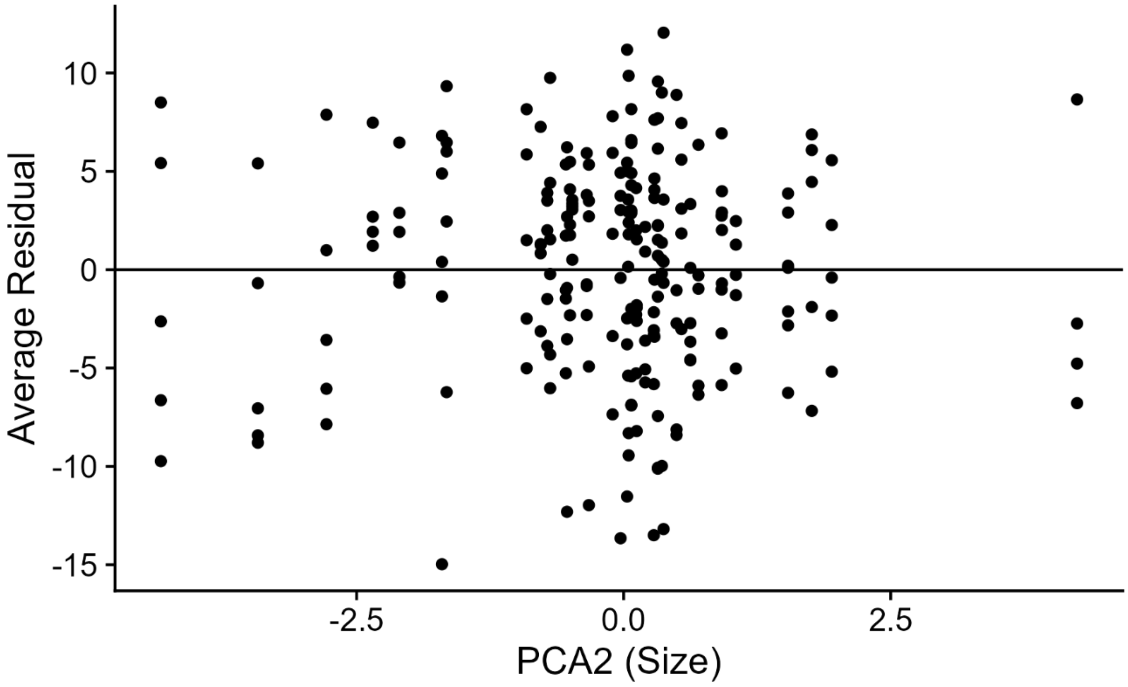


**Figure S7.1.** The average residuals from the posterior of our Bayesian statistical model show no obvious signs of nonlinearity in their relationship with the principal component analysis axis representing *Paramecium* body size (PCA2).

**Literature Cited**

Bürkner, P.-C. (2017). brms: An R Package for Bayesian Multilevel Models Using Stan. *Journal of Statistical Software*, *80*(1), Article 1. https://doi.org/10.18637/jss.v080.i01

Pennekamp, F., Schtickzelle, N., & Petchey, O. L. (2015). BEMOVI, software for extracting behavior and morphology from videos, illustrated with analyses of microbes. *Ecology and Evolution*, *5*(13), 2584–2595. https://doi.org/10.1002/ece3.1529

R Core Team. (2023). *R: A Language and Environment for Statistical Computing*. R Foundation for Statistical Computing. https://www.R-project.org/
